# A personalized eye-tracking system reveals complex state-dependent visual acquisition in lizards

**DOI:** 10.64898/2026.01.06.697876

**Authors:** Nimrod Leberstein, Tali Remenick, Milan Becker, Mark Shein-Idelson

## Abstract

Gaze shifts are fundamental to visual exploration and processing, yet empirical knowledge is largely confined to a small number of model species, limiting our understanding of general principles of visual acquisition across animals. Here, we introduce a versatile, low-cost and open-source framework design for eye tracking in small freely-moving animals. Leveraging photogrammetry, 3D modeling, and 3D-printing, we produce headmounts adaptable to diverse head-eye geometries as we demonstrate for lizards, turtles, and mice. Detailed recordings in Pogona vitticeps, a laterally eyed ectotherm with a high-acuity retina, reveal a rich oculomotor repertoire with kinematics closely resembling the saccadic main sequence in primates. We found both movement-coupled large binocular saccades which exhibited a strong axial bias as well as small, isotropic, and monocular saccades suggesting a capacity for visual exploration of local details. Visual acquisition strategies were strongly tied to behavioral states with periods of active sampling (3-5Hz saccades) separated by periods of ocular stillness lasting many seconds and suggesting divergent visual acquisition strategies for ectotherms. Further, active states were correlated with pupil constriction in contrast to mammals and similarly to birds. Our results demonstrate the utility of our experimental approach for understanding visual acquisition across terrestrial vertebrates. Such a multi-perspective view on natural visual acquisition across species will be essential for developing a generalized understanding of vision, its ecological functions, and its evolutionary trajectory.

## Introduction

Shifting gaze is fundamental to visual exploration and processing [1,2]. Each gaze shift comprises a decision to selectively acquire information from a specific region of the visual environment. In foveated animals [3], this process is particularly precise and informative, offering a unique window into the dynamic prioritization of goals across contexts [4,5]. In most studies, gaze information is acquired under head-restrained conditions [6–10]. This setting facilitates precise gaze measurements in controlled conditions, yielding significant insights into visual processing [11–17]. However, under natural conditions, visual exploration involves a dynamic coordination between eye, head and body movements, which is disrupted under head fixation [18–20]. This discrepancy limits our understanding of visual acquisition strategies.

Over the last two decades, new head-mounted systems enabled eye tracking in head-free conditions for humans and primates [21]. These technologies provided valuable insights into the integration of eye movements with broader behavioral patterns [22], such as describing a tight predictive coupling between gaze and gait in humans [23], or showing that hippocampal activity is highly tuned to 3D head orientations and eye movements in macaques [24]. However, these systems are tailored to large heads and not readily compatible for use in smaller model species. Recently, innovative miniature video- and magnetic-based systems were developed, substantially expanding our access to eye measurements in freely moving small animals [19,25–34]. The first video implementation allowed oculography in freely-moving rats through bespoke camera and support modules [25], whereas later mouse-oriented systems further miniaturized the hardware and, in some cases, provided open-source solutions, improving accessibility [26,27,30]. These were complemented by miniaturized magnet-sensor trackers that brought magnetic eye tracking into freely moving small animals without field-bound search coils [28,29]. Still, these valuable systems were typically tailored for a particular species, and rely on non-flexible anchor positions, often tied to chronic electrophysiology or magnetic implants [25–28,30–32,35,36]. Consequently, these did not offer an accessible route for repositioning the optical modules in species with markedly different eye-head geometries, such as broad-skulled, highly lateral-eyed lizards, or head-retracting turtles [37]. This has constrained comparative analyses of visual acquisition across taxa, which is crucial for gaining an evolutionary understanding [38,39].

A particularly important stage in the evolution of visual acquisition is the transition to terrestriality. This transition was accompanied by a major expansion in the potential visual range. Correspondingly, vertebrates evolved larger eyes with higher acuity [40,41]. Following the emergence of amniotes, vertebrates evolved a distinct neck and the atlas-axis bone complex which allowed flexible head movements dissociated from body movement, increased visual sampling range, and improved stabilization during self movement [42]. By contrast, the available evidence from amphibians indicates minimal eye movements, and increased reliance on full body turns for visual orienting [43]. However, the evolution of neural machinery for sampling complex terrestrial visual environments remains uncharted [38,39,44]. Reptiles, the sister class to mammals within amniotes provide a critical comparative perspective on this question [45,46].

Multiple reptile species exhibit complex visual behaviors and possess steep-walled foveae and visual streaks associated with high spatial acuity [47,48]. A limited number of studies in lizards examined visual behaviors. These focused on optokinetic reflexes and prey orientation, without recording eye-movement or performing visual sampling analyses [49–51]. The only quantitative eye-movement datasets in non-avian reptiles are from chameleons. Magnetic eye tracking revealed a high oculomotor range, a monocular eye coordination regime that changes with behavioral context, and a linear relationship between saccade amplitude and speed during prey capture [52,53]. Thus, the spatial and temporal characteristics of visual sampling in non-avian reptiles remain largely unknown [52,54].

To bridge this gap, we developed a flexible system for measuring eye movements in freely moving small animals. Inspired by the use of open-source components for miniaturized video acquisition of the eyes [26], we developed an optical system and a personalization approach which allows positioning the optic modules for diverse head-eye geometries in a variety of animals. Using photogrammetry, we generated 3D-printed headmounts that can be adapted to specific species and subjects, ensuring a precise and stable fit while maintaining minimal interference with natural behavior. We first demonstrate the system’s versatility by acquiring eye video in the lizard *Pogona vitticeps*, the turtle *Trachemys scripta elegans*, and *Mus musculus*. Next, focusing on *P. vitticeps*, a foveated visually orienting, laterally eyed agamid lizard [47,55], we address key questions in visual acquisition. We ask whether agamid lizards exhibit goal-directed ocular saccades. We examine whether visual acquisition strategies in ectotherms differ from those in endotherms across long time scales. We probe the relation between pupil diameter and behavioral states. Studying these questions revealed a complex pattern of visual gaze shifts and lateralized behaviors, as well as state-dependent changes in visual acquisition and pupil diameter in *P. vitticeps*.

## Results

### System design, fitting, and adaptability

To monitor eye movements in freely behaving small animals, we developed a flexible, open-source, low-cost, eye-tracking approach that is applicable across species (Methods, Table 1). A major challenge in implementing eye tracking systems across individuals and species is posed by the diversity in head-eye geometries. To address this, we designed personalized headmounts that are 3D printed based on a 3D model of the animal’s head and eyes as scanned using photogrammetry (Fig. 1a, b). Importantly, this scan followed a surgical implantation of a single hexagonal screw anchor point for connecting the headmount. The model is used to define the desired camera-eye geometry in virtual space (via Fusion360), and produce a single-piece headmount to connect with the anchor, support cameras, and the infrared (IR) mirrors reflecting the eyes (Fig. 1b) (Methods). The headmount is 3D printed as a single piece (without the mirror holders which are printed separately with interlocking connectors). When a photogrammetry scan is not an option, the modules can be printed as individual interlocking pieces which form a modular solution kit, such that the fitting procedure can be performed directly on the animal (Fig. 1c) (Methods). To achieve sharp images under shorter working distances (1.5–5cm) without background contamination, stock camera optics were replaced with manual focus lenses fitted with an IR-pass filter (Fig. 1d), and two IR LEDs were wired and glued to each camera module aimed at the mirrors [26]. This provided stable illumination of the eye and synchronization verification via programmed power-off windows (Methods). The complete headmount with LEDs, wires, and cameras (2.13g each) weighed between 8-10g depending on animal size. Before recording, headmounts were connected to the animal and counterbalanced by a pulley system holding the headmount and connecting cables. To assess the stability of the system, we quantified camera jitter (spatial cross-correlation peak shift between the first frame and other frames calculated on a stationary image segment) during free behavior. The typical measured jitter was small (0–100µm) (Fig. 1e) and could be corrected by subtracting the jitter amplitude.

**Figure 1:**
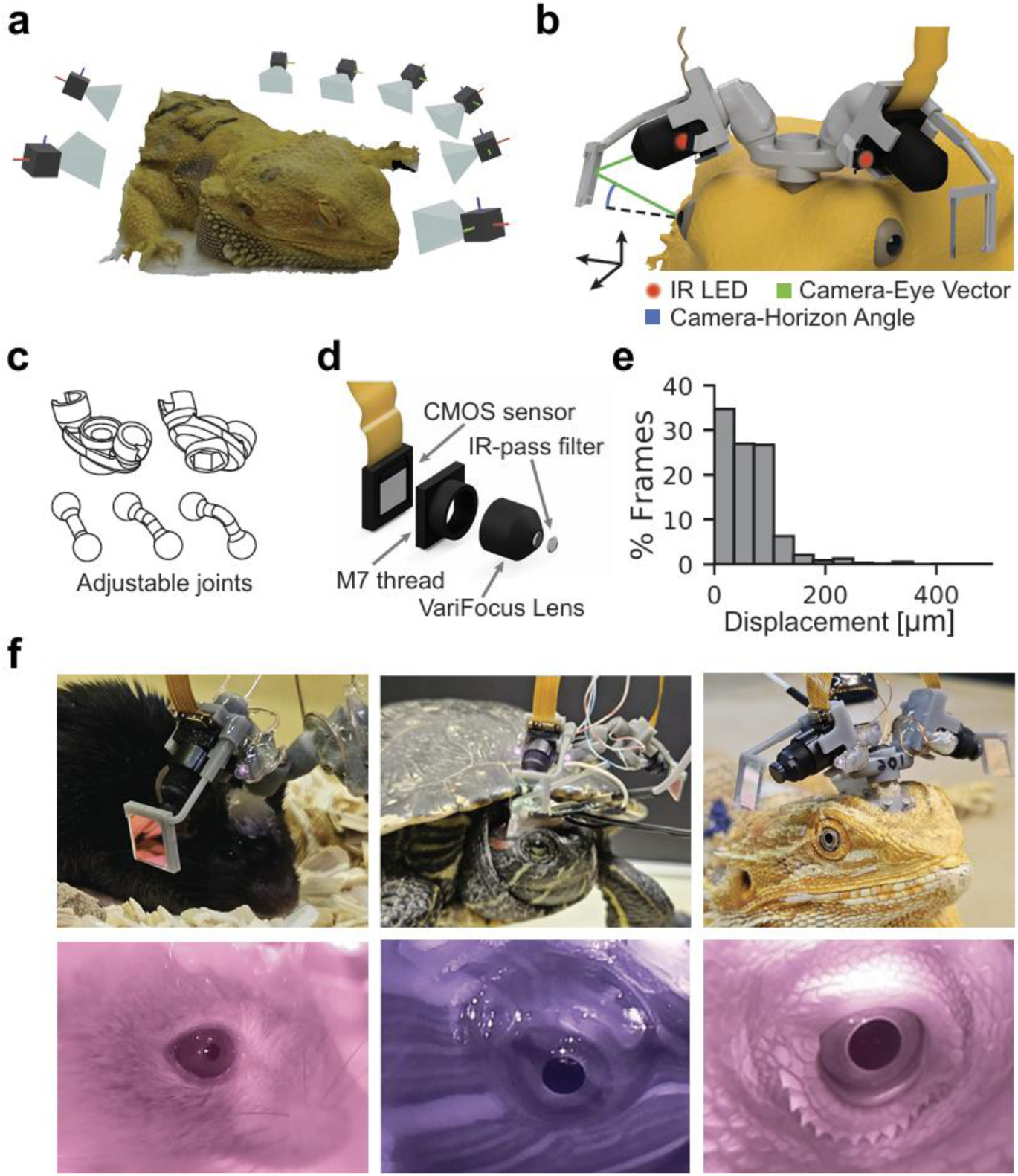
A model-based approach for 3D printing personalized headmounts for eye tracking in small freely behaving animals. **a.** 3D animal model acquired using photogrammetry by combining multiple camera viewing angles (a subset shown for illustration) **b.** A lizard’s head (3D rendered) fitted with a full single-print camera headmount (gray). A virtual fitting procedure is used to calculate the camera (black) and mirror position such that a light beam (green line) from the camera center reaches the eye center. The inclination angle is determined by an animal-centric horizon defined as a line connecting the two eye centers (dashed black). Inclination angles fluctuate between 20-30 degrees for different animals. After 3D printing the headmount, two IR LEDs (red) are glued to each camera holder to provide illumination at 940nm, and an IR reflector is glued to an alignment groove on the mirror holder. **c.** Diagrams for a modular solution kit. Different ball-joint arms allow a flexible and adjustable fit for variable head geometries without the need for 3D scanning. **d.** The lens assembly of purchased minicameras is removed and replaced by a long M7 thread and 3.7mm manual focus lens to enable variable working distance. The lens is fitted with an IR-pass filter to remove image contamination by visible light. **e.** Histogram of the camera movements, quantified as percentage of overall frames in each bin, indicating low camera jitter (over 90% of frames under 200µm). **f.** Examples of personalized headmounts for a mouse (*Mus musculus*), turtle (*Trachemys scripta elegans*) and lizard (*Pogona vitticeps*) with corresponding raw camera images acquired from the eye.

**Table 1.**
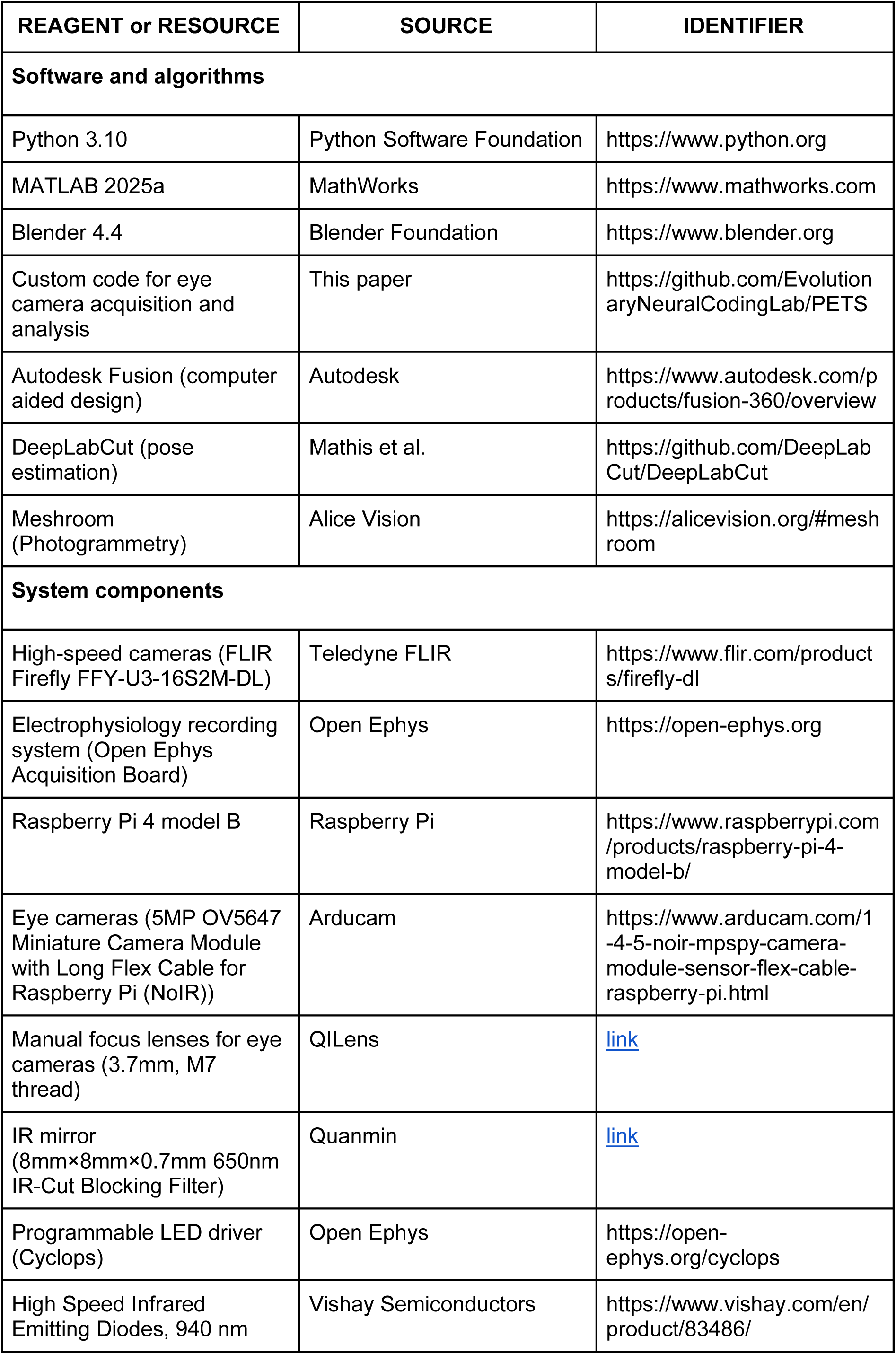
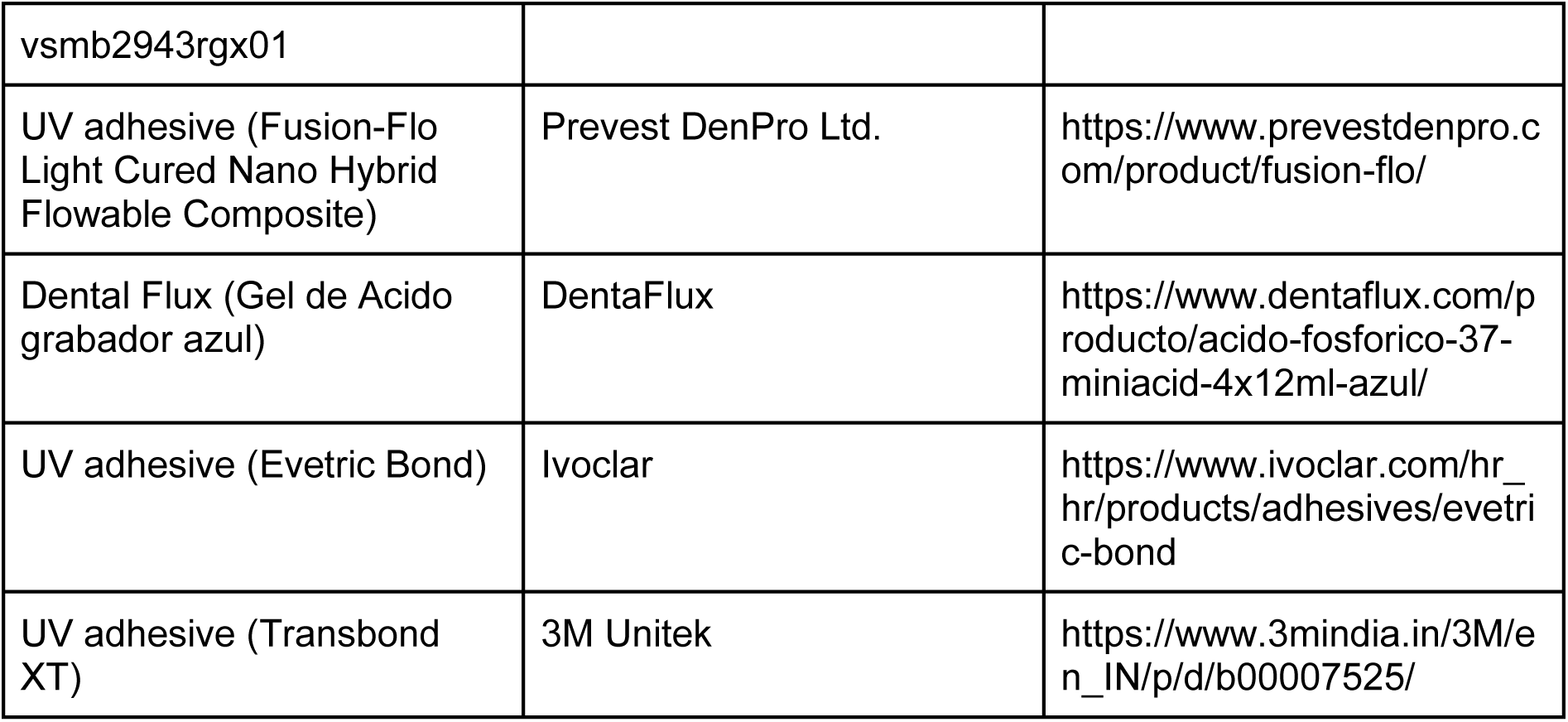
Table of system components and software.

To demonstrate the system’s versatility, we used it to successfully record eye movements in three distinct model animals: bearded dragons (*Pogona vitticeps*), red-eared slider turtles (*Trachemys scripta elegans*), and mice (*Mus musculus*). These species were chosen for their anatomical and behavioral diversity (Fig. 1f): *P. vitticeps* lizards are typified by wide heads and lateralized eyes; turtles posed a unique challenge due to their ability to retract and extend their heads into their shells; and mice are small and abundant in research. To accommodate measurement in turtles, we incorporated an extender between the anchor point and the headmount connector, allowing full range of head movement by keeping the headmount outside the shell when the head is retracted [56]. The longer working distance between the eyes and cameras was compensated by the manual focus of the modified lenses. Mice required a reduced anchor size and headmount design to account for their smaller size. To accommodate this, the hexagonal brass threaded spacer (0.5 grams) was replaced with a lighter plastic version, and the headmount design was scaled down to a total of under 5g.

### Eye movements repertoire in freely behaving *P. vitticeps*

The intricate musculature and neural circuitry controlling vertebrate eye movements enable diverse oculomotor behaviors that reflect visual acquisition strategies [2]. Yet, limited measurements in freely moving conditions and across species hinder the derivation of general principles [7]. To study the oculomotor behavior in *P. vitticeps* – a visually orienting agamid lizard with highly lateralized eyes – we acquired bilateral eye videos from freely exploring animals (n=5) placed in a behavioral arena (Video S1) [57,58]. Eye videos were used to reconstruct egocentric gaze vectors (eye-in-head angles) from pupil annotations using a weak perspective model (Fig. 2a) [59,25,60,36,35,33], validated using a 3D Blender-based [61] simulation of the eye-camera system (Methods, Fig. S1).

**Figure 2:**
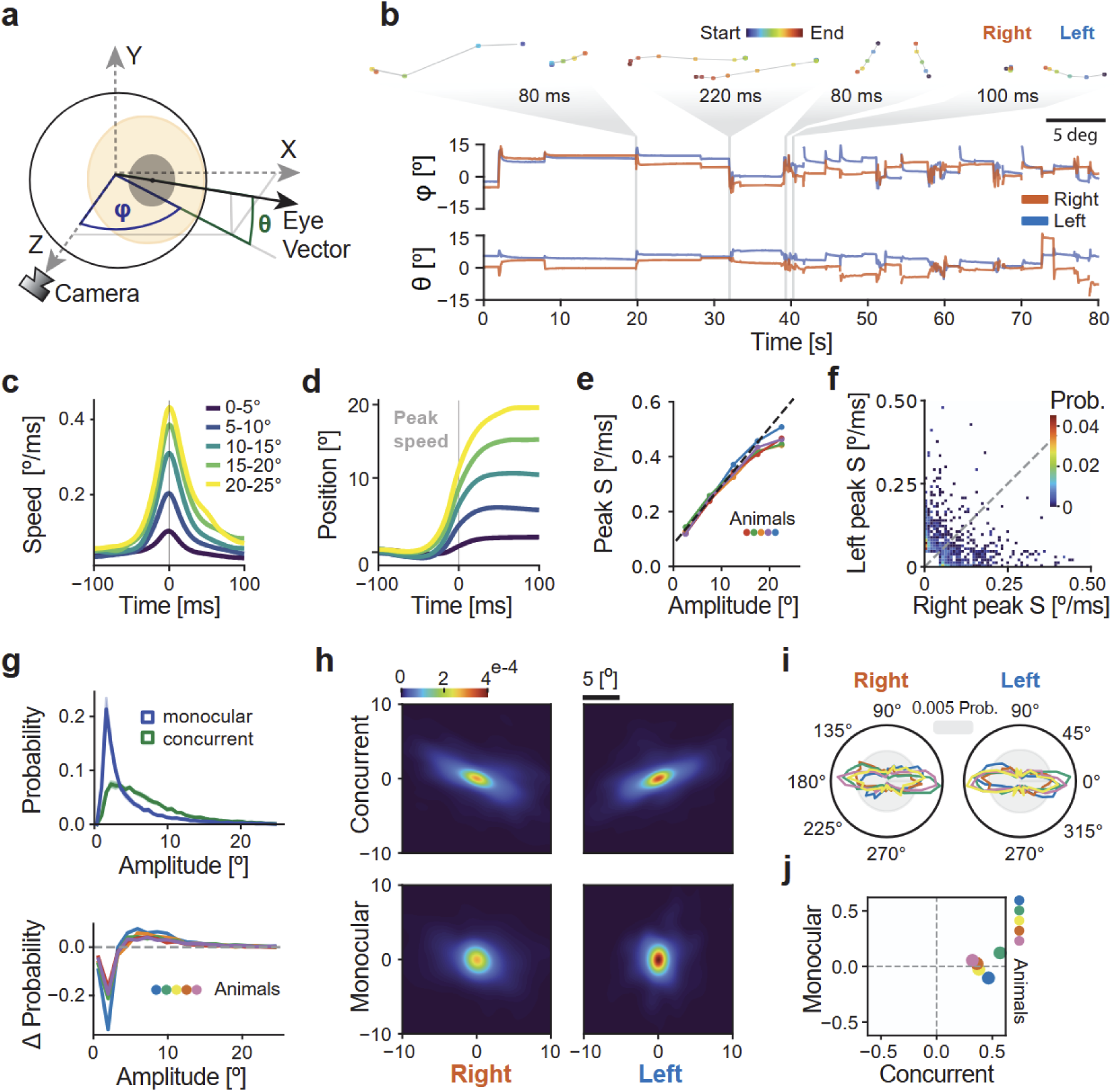
Main saccadic sequences with isotropic monocular and directionally biased concurrent saccades in *P. vitticeps*. **a.** Schematic of calculating viewing angle in camera coordinates. 2D data points from eye videos were converted to 3D directions defined as azimuth (φ) and elevation (θ) angles (Methods). **b.** Simultaneous eye angle tracking for both eyes ((φ,θ)=(0,0) is the eye’s median position). (Top) The spatial trajectory of four example saccades of different types. left to right: divergent, conjugated, disjunct and monocular dynamics. **c.** Mean saccade angular speed profiles for one animal (n=6954 saccades). Events were binned by net amplitude (5° bins, color coded), aligned to t=0 ms (peak speed), and smoothed with a 10 ms gaussian kernel. **d.** Mean angular-displacement traces (from (c)) along each saccade’s movement axis, aligned to onset. **e.** Relationship between saccade amplitude and peak angular speed. Each colored line represents one animal (n = 5; 34,876 saccades). Linear fit y = 0.021 [ms⁻¹] × amplitude [°] + 0.083 [°/ms] (R^2^ = 0.44, r= 0.66, p<5e-10). **f.** Inter-ocular peak-speed coupling. Two dimensional probability histogram of right- vs left-eye peak angular speed [°/ms] for all saccades during head-stationary epochs pooled across animals (equal-animal weighting). **g**. (Top) Distribution of saccade amplitudes for concurrent (green) and monocular (blue) events. Mean (±S.E.M shading) animal-wise probability distributions (0-25°). (Bottom), Per-animal difference curves (concurrent - monocular). **h.** Heatmaps of eye-endpoint densities for concurrent and monocular saccades (single animal) relative to saccade start location, plotted in degrees on the unrotated camera axes. **i.** Polar distributions of saccade directions per eye and animal, rotated to align the eye’s dominant axis of motion to the 0/180 degrees axis (light gray circle= 0.005 density). **j.** Axial bias [A.U.] of saccade directions comparing the axial bias of concurrent (x-axis) and monocular (y-axis) saccades (animals color coded). For each dataset: bias = (P(main-axis) - P(perpendicular)) / (P(main-axis) + P(perpendicular)).

Our recordings revealed that freely behaving *P. vitticeps* express a diverse oculomotor repertoire, with movements varying in speed, conjugacy, and temporal density over the course of each session (Fig. 2b). Across animals, gaze vectors covered an average range of 35.0° ± 7.3° along the main movement axis (defined as the orientation of maximal bidirectional saccadic density) and 26.1° ± 6.4° along the perpendicular axis (Fig. S2). These spans were measured relative to each eye’s resting orientation, which varied modestly between recordings (σ_main-axis_=4.8°, σ_perpendicular_=2.9°). Within this workspace, gaze redirections often occurred as brief, discrete high-speed events, as next analyzed.

Saccades are rapid eye movements that abruptly shift gaze, minimizing transit time while maximizing accuracy [62,63]. To characterize the kinematics of saccadic eye movements in *P. vitticeps*, we identified saccades using an angular speed threshold of 50°/s applied to gaze-angle traces. We detected multiple binocular configurations, including conjugate, divergent, convergent, disjunct (unequal inter-ocular dynamics), and monocular events (Fig. 2b). Saccades exhibited stereotyped speed profiles characterized by rapid acceleration to a single peak, followed by smooth deceleration back to baseline (Fig. 2c). The corresponding displacement showed a stereotyped sigmoid-shaped transition from the pre-movement baseline to a new gaze position, with most of the change occurring over a brief interval, and with a shallower tail as the movement approached its endpoint (Fig. 2d). Across animals, peak angular speed scaled systematically with saccade amplitude, forming a linear relationship across most of the measured range, with a tendency toward saturation at higher amplitudes (>20°) (Fig. 2e).

In lateral-eyed rodents such as mice, eye movements during free behavior are tightly coupled with head-led gaze shifts, supporting faster transitions and providing image stabilization [36]. In such an acquisition scheme, the two eyes rotate in temporal coordination, with no monocular saccades. To quantify inter-ocular coordination in *P. vitticeps*, we distinguished between concurrent saccades (binocularly synchronous, but not necessarily directionally conjugated), in which both eyes crossed the speed threshold within a narrow temporal window (<34ms), and monocular saccades, detected in only one eye. We found monocular (highly asymmetric) saccades in all animals and across all recording sessions (Fig. 2f, S3, Video S2). We next compared the amplitude distributions between monocular and concurrent saccades. While all saccades were biased towards smaller amplitudes, monocular saccades were typically smaller than concurrent events (Fig. 2g). Thus, while the head is stationary, monocular saccades are fast and local while concurrent saccades are larger, possibly reflecting gaze shifts during locomotion or attention shifts rather than local detail extraction.

To support the interpretation that some saccades reflect active gaze redirection rather than purely vestibulo-ocular stabilization, we asked whether they could be observed when the head is still. We recorded head kinematics using a head-mounted accelerometer and identified no-movement epochs (Methods). While most saccades occurred concomitant with head movement, a considerable fraction were detected during head-stationary periods (33.2 ± 9.4%, mean ± SEM; n = 15,613 events, 4 animals). Further, monocular saccades constituted almost half of all head stationary events (48.9 ± 8.8%), whereas head-led saccadic events were less frequently monocular (21.5 ± 5.5%). Correspondingly, there was a small but significant correlation between saccade amplitude and movement (Monte-Carlo shuffle (n = 10,000): Δ = 0.64° (n = 4), p = 1×10⁻⁴). Thus, while most gaze shifts occur during movement and play a role in gaze stabilization, a considerable fraction of eye-led monocular (and binocular) gaze shifts occur during stillness, suggesting a shift in visual attention.

During free exploration, lizards frequently traversed the arena and produced repeated gaze shifts as they moved. Because directionality biases in eye movements often mirror ecological demands on visual sampling [2], we asked whether saccades exhibit structured directional organization. To test this, we expressed each event as a displacement vector in eye-centered coordinates and aligned all saccades to a common origin. We found that for concurrent saccades, relative endpoints clustered strongly along a single dominant axis, revealing a pronounced axial organization of gaze displacements. We note that these axes are tilted because cameras are aligned to the head-to-tail axis, while animals usually hold their heads at an inclined angle to this axis. We therefore rotated distributions to align to the main axis and revealed that this bias for concurrent saccades was consistent across animals (Fig. 2i, j). In contrast, monocular saccades exhibited a broader and more isotropic distribution of endpoints (Fig. 2h, j). In summary, we find a continuum of saccade profiles that shift between two separate modes of visual acquisition. During animal movement we find large, concurrent eye movements with a clear directional bias which can partially be explained by gaze stabilization. During stillness, when gaze stabilization is not active, we find smaller, isotropic eye movements that shift focus to small sub regions in the visual environment.

### Behavioral state affects ocular temporal dynamics and pupil dilation in *P. vitticeps*

In endotherms, spontaneous eye movements during wakefulness occur at a relatively stable rate of ∼ 3-4 Hz [2,62]. As ectotherms, reptiles may employ a different temporal visual acquisition strategy shaped by their metabolic constraints and energy conservation demands [64]. To investigate the temporal dynamics of visual acquisition, we monitored eye and head movements for extended time periods (2-3 hours). We observed that eye movements shifted between a state with a limited number of eye movements (Fig. 3a) and a state with a high frequency of eye movements (Fig. 3b). To examine if these visual acquisition states coincided with animal movements, we segmented the behavioral recordings into Active states (during locomotion and head movements) and Quiet epochs (periods of stillness) using concurrent accelerometer measurements (Methods). Active states constituted 10.2% of the total analyzed recording time. These epochs were accompanied by frequent eye movements (3-5 Hz, Fig. 3b,c). In contrast, during Quiet states saccade frequency dropped under 1Hz (Fig. 3a,c). This effect was significant (Monte-Carlo shuffle (n=10,000): Δrate = 2.66 Hz, p = 1×10⁻⁴). Notably, prolonged quiescent periods, devoid of any detectable eye movements, lasting up to several tens of seconds, were common (Fig. 3a, d), suggesting discrete shifts between exploratory and stationary visual states. While these long intervals were widely distributed, an increased probability of 1-3s intervals was observed across animals suggesting a typical time scale during Quiet states (Fig. 3d). Thus, *P. vitticeps* exhibits a transition between active and quiet behavioral modes reflected in both movement and visual acquisition.

**Figure 3:**
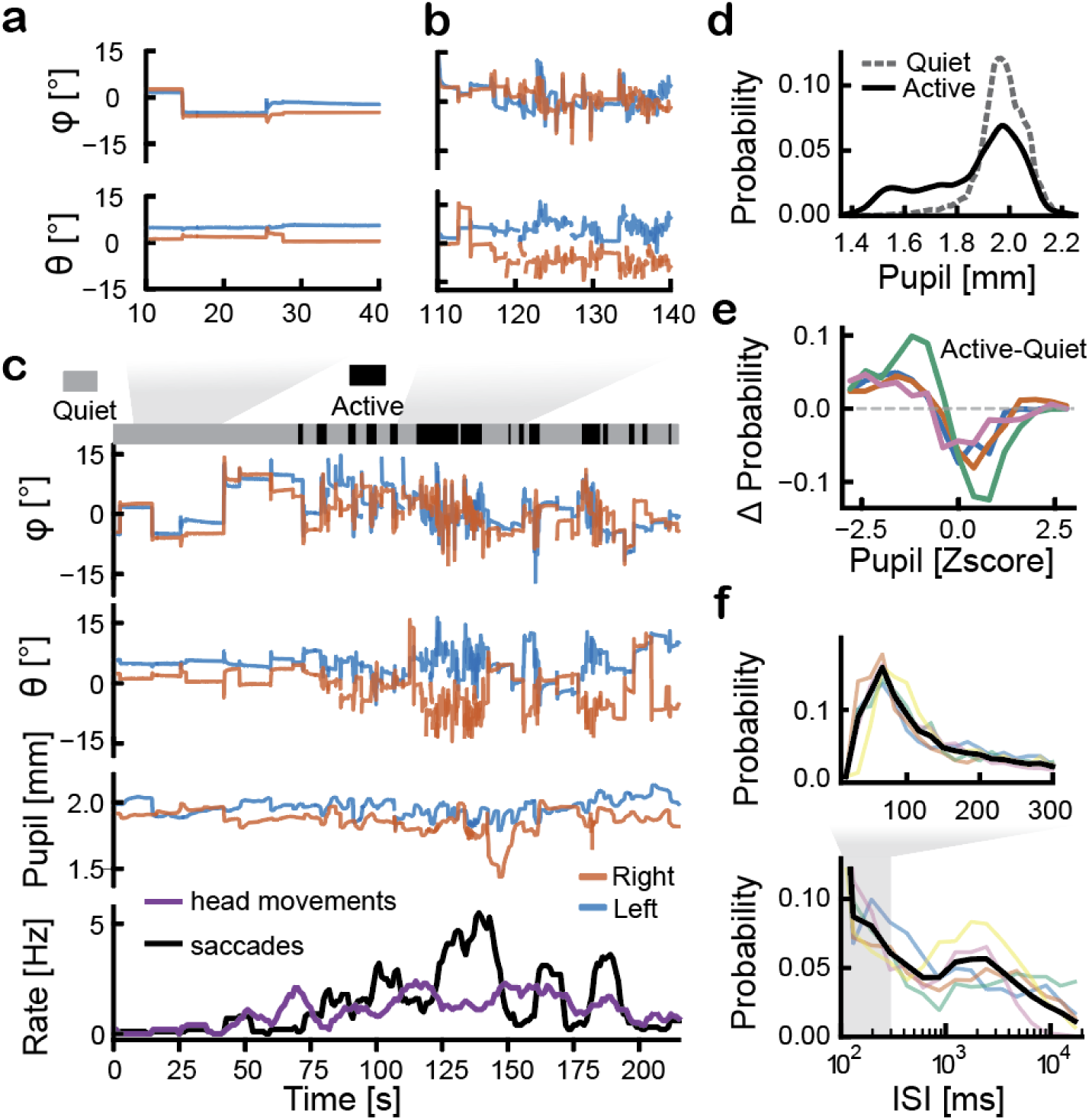
Visual acquisition profiles change with behavioral state and pupil diameter. **a.** Eye angle dynamics for 30 seconds during a behaviorally quiet epoch. Angular traces were median-centered to each eye’s resting position. **b.** As in (a) but for an active period. Note frequent saccades. **c.** Eye angles, pupil size, and rate of saccade and head movement events acquired during free behavior. The top bar indicates behavioral classification into active (dark) and quiet (light) states with typical epochs enlarged in (a) and (b). Head movements (purple) and binocular saccade rates (black) were computed in sliding 10s windows (1s step). **d.** Inter-saccade interval (ISI) distributions. ISI probability histograms shown on linear (top) and logarithmic (bottom) scales, with overlapping domains between 200-400ms. Each trace represents one animal with the average over animals in black. **e.** Pupil diameter probability distribution for active (solid) and quiet (dashed) behavioral states for one animal (both eyes). **f.** The difference between z-scored (mean/std) pupil-diameter distributions in active and quiet epochs. Positive deflections indicate pupil sizes that occur more frequently during Active periods.

The changes in behavioral state prompted us to examine the dynamics of pupil diameter. In mammals, pupil diameter is often binocularly coupled, and scales positively with arousal, providing an external index for cognitive effort and attention [65–68]. To discern inter-ocular pupil behavior in *P. vitticeps*, we computed left-vs-right pupil correlation values (Methods). Pupils showed clear positive correlations during 66.3 ± 7.9% of the time (Pearson r > 0.2) (Fig. 3c, S4). Whereas pupils were uncorrelated for 23.4 ± 3.5% of the time (−0.2 ≤ Pearson r ≤ 0.2) and anti-correlated for 10.3 ± 4.9% (Pearson r < -0.2). Despite this divergence and given the transitions in behavioral states (Fig. 3d-e), we wanted to understand if overall pupil diameter adheres to some relationship with arousal. To test state-dependent modulation, we compared pupil diameter distributions across behavioral states over long recording sessions (9.2 hours over 5 animals) (Fig. 3c, e). The long recording periods and uniformly illuminated arena were instrumental at averaging out illumination-specific effects. We found a significant difference (Monte Carlo shuffle, n=20000 p = 5.0e-05) between states: pupils were smaller during Active epochs and dilated during Quiet epochs (Fig. 3e). This difference was consistent over animals (Fig. 3f). Interestingly, it is inverse in directionality to the one found in mammals.

## Discussion

We developed a versatile and technically accessible approach for eye tracking in freely moving small animals. By combining 3D photogrammetry with personalized 3D-printed headmounts, we established a framework that is rapidly adaptable across species, as demonstrated across three species with distinct cranial morphologies (Figure 1). Focusing on *P. vitticeps*, we found a rich oculomotor repertoire comprising gaze stabilizing saccades during movements with axial biases, and exploratory monocular and binocular saccades without movements (Fig. 2). Finally, we found that saccades dynamically changed throughout long recordings, exposing a bi-phasic visual acquisition strategy that was tied to behavioral state and pupil diameter fluctuations (Fig. 3). The implications of these results on our understanding of visual acquisition strategies across species are discussed below.

Visual strategies across species depend on the degree of head motility, the position of eyes, and the morphology of retinas, and are reflected in saccade kinematics and oculomotor movement statistics [69]. We found that eye movement spans are anisotropic with a clear bias to larger movements in one orientation (Fig. 2h-j). Comparable directional anisotropies have been reported in other amniotes, including primates, rodents, chameleons and birds [29,53,70]. Such anisotropy likely reflects a combination of biomechanical constraints of the oculomotor plant [71,72] and the nonuniform distribution of behaviorally relevant visual targets across the sampled environments [73]. Another property with striking similarity to visually oriented amniotes is saccade kinematics [2,62]. Individual eye movements in *P. vitticeps* showed the characteristic rapid acceleration and deceleration phases that define vertebrate saccades [38,74] (Fig. 2c, d), and their peak speed scaled linearly with amplitude (Fig. 2e). This relationship largely resembles the ’main sequence’ previously described in primates and cats [75–77] with a similar rate slope of ∼20 [1/s] in both primates [77] and *P. vitticeps* (Fig. 2e). Interestingly, the main sequence curve differed at high saccade amplitudes for which primates exhibit strong saturation in contrast to only modest saturation in *P. vitticeps*. This could result from factors such as eye geometry. However, the similarity in the saturation value for both primates and *P.vitticeps* (∼15°) suggests that this discrepancy results from the larger oculomotor span (90° [2,78] in humans vs 35° in P. vittices, Fig. S5). Indeed, the absence of individual eye movements larger than 25° in *P. vitticeps* likely reflects the animal’s reliance on head movement contributions during broader gaze shifts, confining saccades to the low-amplitude regime where peak speed is not yet saturated [63]. In that sense, the non-saturated profiles in *P. vitticeps*, and the reliance on head movements (∼40% of gaze shifts were executed when the head was stationary, whereas the remaining ∼60% occurred during coordinated eye-head saccades) are advantageous relative to the saturated profiles in humans. This may reflect a limited ability to move large heads fast enough. Alternatively, the estimates of the natural span in humans (and the consequent speed saturation with amplitude) may be biased by the head-restraint conditions used in many experiments [75,77,78]. This is consistent with the lack of large ocular excursions during naturalistic, head-free behavior [77,79,80].

Across taxa, saccades can serve different functional roles, ranging from compensatory quick phases that accompany head movements to voluntary shifts that reposition the eye toward behaviorally relevant targets [2]. In many lateral-eyed mammals such as rodents and ungulates, saccades are reported primarily as quick phases supporting head-led gaze shifts and vestibular correction [36,38]. Such animals rely predominantly on head rotations to redirect gaze with minor oculomotor span (∼20° for mice [36]). This behavior fits the lack of a clear localized fovea or the existence of a visual streak [81–83]. By contrast, *P. vitticeps* frequently generated saccades in the absence of head movements, consistent with eye-led reorienting similarly to primates. These events likely reflect goal-directed ocular shifts, and are supported by evidence for a fovea in agamid lizards [47]. We note however, that foveated animals can perform goal-directed visual behaviors also using head movements. Avian reptiles can quickly direct gaze using their heads, with some species exhibiting only minimal eye movements (∼3° for barn owls [84,85] and ∼5° in chickadees [86]) while relying on head movements for both saccades and image stabilization during locomotion [3,87,88]. In contrast, the propensity for eye-only saccades in *P. vitticeps* is reminiscent of some acquisition strategies in foveated primates and head-orienting birds. In primates, the existence of directed, time-stamped saccades have been widely leveraged to study attention and decision-making [5,89]. The presence of similar eye-led shifts in *P. vitticeps* suggests a tractable route for extending visual attention studies to reptiles, thus providing a comparative perspective that can shed light on the evolution of attention and corresponding neural processing [90].

In laterally eyed species, monocular saccades can be advantageous because portions of the visual field are accessible to only one eye, making unilateral shifts a direct and efficient means of sampling the most relevant features of each hemifield [48,91]. However, this potential is not uniformly expressed across taxa: many lateral-eyed mammals and birds exhibit tightly yoked ocular rotations [29,36]. Against this backdrop, we observed flexible ocular coupling in *P. vitticeps* (Fig. 2a, f, g). Asymmetric and monocular saccades were common, forming a graded continuum of coupling strengths (Fig. 2f). This regime is consistent with proposals that monocular and conjugate saccades may arise from the same control architecture, with coupling strength being a tunable parameter [91]. In this view, monocular independence and binocular conjugacy represent endpoints of a shared control space, with coordination varying by context. Chameleons, for example, exhibit extreme ocular independence to support crypsis during scanning, yet transiently increase coupling for prey capture [52,92]. By contrast, mammals with front-facing eyes, including primates and several predatory rodents, exhibit reinforced binocular coupling that favors tightly yoked rotations for stereopsis and coordinated pursuit [33,93]. *P. vitticeps* exemplifies a balanced regime in which modest monocular independence coexists with reliable binocular coordination.

The temporal structure of eye movements in *P. vitticeps* revealed a striking departure from patterns described in endothermic vertebrates. Lizards often remained without ocular motion for several tens of seconds, intervals far longer than the continuous high-frequency sampling typical of endothermic vertebrates, including birds [94], rodents [36,95], and primates [16,62,96,97]. In *P. vitticeps*, these prolonged quiet states alternated with relatively brief epochs of dense saccadic activity (active states), producing a bimodal distribution of inter-saccadic intervals (Fig. 3). This suggests that reptiles may alternate between two distinct visual acquisition modes which are presumably more energy efficient. This behavior aligns with the well-documented postural immobility in ectotherms [98], but the accompanying stillness of the eyes appears distinctive and indicates a state in which visual sampling demands differ markedly from those that drive active scanning [99]. How the brain uses these quiet periods to process information remains to be studied [100,101].

In mammals, behavioral states are linked with brain states and pupil diameter across both sleep and wakefulness [36,65–68,95,102]. We found pupil diameter in *P. vitticeps* constricted during active, movement-rich states and dilated during quiet wakefulness. This polarity contrasts with the dilation-with-arousal pattern characteristic of mammals [66–68]. A similar inversion has recently been documented in birds, where pupil constriction accompanies aroused wakefulness and REM-like sleep [103]. Because both reptiles and birds possess predominantly striated iris musculature [104], this cross-taxon correspondence suggests that rapid, behavior-linked pupil adjustments may reflect a broader sauropsid pattern rather than a feature unique to birds. Since the functional basis for pupillary state-dependence remains unclear, the existence of highly structured sleep state transitions in *P. vitticeps* [105] could provide valuable clues for its function.

Taken together, our findings highlight the value of comparative head-free eye tracking approaches. Recent technological advances provide unprecedented access to structural and functional properties of eyes and retinas on both the cellular and molecular level and across species [106–109,69,110,111]. Our work complements these experimental advances by enabling precise quantification of ocular behavior in species that have been largely inaccessible using conventional eye-tracking techniques. Going forward, the compatibility of our method with electrophysiology and multi-camera tracking provides a practical route for extending our understanding of visual neural processing across taxa. Our results demonstrate how expanding eye-tracking to reptiles can facilitate the extraction of conserved acquisition strategies (e.g. the saccadic main sequence and axial biases in scanning) as well as specializations (e.g. ectothermic acquisition modes and modulation of pupil diameter by behavioral state). Such a multi-perspective view on natural visual acquisition across species will be essential for developing an understanding of vision, its ecological functions, and its evolutionary trajectory.

## Data and code availability

The source data to produce the figures, code used to collect and analyze the data, 3D models, virtual fitting environment, preprocessing pipelines and step-by-step instructions are available at https://github.com/EvolutionaryNeuralCodingLab/PETS

Any additional data is available from the corresponding author upon a reasonable request.

## Acknowledgements

This project has received funding from the European Research Council (ERC) under the European Union’s Horizon 2020 research and innovation programme (Grant agreement No. 949838, MSI) and from the Israel Science Foundation (ISF, grant No. 1597/25, MSI). The authors are most grateful to A. Shvartsman for technical and administrative assistance; the animal caretaker crew for lizard care; N. Albeck for comments on the manuscript; G. Gorfung for assistance with eye vector calculations; the Shein-Idelson laboratory for their suggestions during this work.

## Ethics Declarations

The authors declare no competing interests.

## Contributions

M.S.I. and N.L. initiated and designed the project, N.L. and M.B collected the data, N.L. and T. R. analyzed the data, N.L. performed the statistical analysis, N.L. and M.S.I wrote the paper. M.S.I acquired the funding.

## Materials and methods

### Animals

Eye movements were recorded from five dragon lizards (*Pogona vitticeps)*, two red-eared slider turtles (*Trachemys scripta elegans)*, and one mouse (*Mus musculus)*. Lizards were obtained from commercial suppliers and housed in a dedicated animal facility at the I. Meier Segals Garden for Zoological Research at Tel Aviv University, under a 12:12 hour light-dark cycle (07:00-19:00) at 24°C. Lizards were fed mealworms (*Tenebrio molitor*), cockroaches (*Nauphoeta cinerea*), and vegetables, with water available ad libitum. Red-eared slider turtles were maintained in an enclosed outdoor pond at the zoological garden of Tel Aviv University (Approval number: H10900/2024). Before experiments, turtles weighing between 800 and 2000 grams were transferred to an indoor laboratory tank (100×120×74 cm; W×L×H) equipped with a basking platform and heat and light sources, under a 12:12 hour light-dark cycle (07:00-19:00) at 24°C. Male and female mice (C57BL/6) were used. Animals were housed under standard vivarium conditions (22°C, 12-h light/dark cycle, with ad libitum food and water). All experimental protocols complied with relevant ethical regulations for animal use and were approved by the institutional animal care and use committee at Tel Aviv University (Approval numbers: TAU-LS-IL-2410-145-4, TAU-LS-IL-2411-147-4).

### Eye tracking system

Eye movements were recorded using a custom dual-camera IR eye-tracking system. Two miniature IR-sensitive cameras (NoIR Spy Camera for Raspberry Pi, Arducam OV5647 sensor) each controlled by an independent Raspberry Pi 4B. Cameras were aimed at the eyes via small (8×8mm) 940nm IR-reflective mirrors connected to a personalized, single-piece 3D-printed headmount. The headmount was designed from a photogrammetry-derived 3D model of each animal to define precise camera-eye geometry and ensure unobstructed visibility of both pupils. Mirrors were held in separately printed holders, allowing rapid replacement of this delicate component without modifying the rest of the headmount. Each camera was fitted with a QILENS 3.7mm manual-focus M7 lens to account for the short working distances (∼ 1.8-4 cm) and a custom-cut IR-pass filter (high-pass > 850nm). Illumination was provided by two 940 nm IR LEDs per camera, wired through a Cyclops LED driver for precise control and synchronization verification via predefined LED-off events (see Data Synchronization below). To allow rapid, repeatable connection across sessions, the headmount attached to a chronically implanted hexagonal standoff (brass M3 (0.6g) for lizards and turtles, plastic M1 (0.15g) for mice), and the underside of each headmount has a matching hexagonal recess. This keyed interface constrained rotation and ensured that the headmount always returned to the same orientation relative to the eyes. Rarely, animals collided with the arena boundaries, breaking the mirror holders extending from the headmount. We therefore integrated generic holders that interlock to the headmount, restoring the predefined geometry upon replacement.

### Anchor Implantation Surgery

To provide a stable connection point for the headmount, animals were implanted with a lightweight cranial anchor. Surgeries were performed under isoflurane anesthesia (4%) with perioperative analgesia (meloxicam 0.2 mg/kg or carprofen 2 mg/kg) and prophylactic antibiotics (enrofloxacin 5 mg/kg). Following exposure and cleaning of the skull, the hexagonal anchor (brass M3 or plastic M1) was affixed using UV-curing adhesive and dental cement. In reptiles and turtles, where skull thickness permits, 2-4 miniature surgical screws (M1) were inserted to increase long-term stability; mice were implanted without screws due to their thin cranial bone. Ectothermic animals (*P. vitticeps*, *T. scripta elegans*) typically resumed normal activity within ∼12 hours, whereas mice recovered after ∼2 hours. Photogrammetry scans for headmount design were performed under anesthesia once the anchor has fully stabilized (after cement curing, typically within minutes). Detailed surgical steps are provided in Supplementary Methods.

### Turtle-specific adaptations

Because turtles retract their heads during unconstrained behavior, we set the anchor point for the system on an extender module to support the hexagonal anchor. The module length (50mm) was calibrated to allow the animal full head retraction of the head without causing a collision between the carapace and headmount. To provide a connection point for the extender module, an M2 hex nut was connected to the module and embedded in dental cement, creating a detent that enabled a reliable 3D model and precise repositioning between sessions. An early version of the system was used for data collection in Becker et al. [56]

### Photogrammetry-based animal modeling

High-resolution 3D models of each animal’s head were generated to enable precise virtual fitting of the personalized headmount. While animals were under anesthesia, 40-80 photographs were captured from multiple viewpoints using a Sony RX100-II camera under diffuse room lighting, ensuring complete coverage of the head-anchor-eye geometry. Images were processed with the open-source photogrammetry pipeline Meshroom (AliceVision), producing 3D textured mesh of the scene. Scale was set using the known physical dimensions of the implanted standoff, which provided a rigid and easily identifiable reference for the model. Calibrated meshes were inspected for reconstruction artifacts and, when necessary, lightly cleaned (e.g., removal of floating surfaces and background) before imported into CAD software (Fusion360) for headmount design and virtual camera alignment. Full photogrammetry settings and parameters for Meshroom are provided as a project file in the associated repository.

### Headmount virtual fitting and fabrication

Personalized headmounts were designed in Autodesk Fusion 360 using the scaled photogrammetry-derived meshes as geometric reference. Rather than sculpting full headmounts de novo, we assembled each design from a library of standardized functional modules (camera holders, mirror holders, and anchor interfaces) that were positioned around the 3D head model to define the desired camera-eye geometry. These modules were then joined into a single rigid piece using simple lofts and connectors, yielding a reproducible headmount that preserved the designed spatial relationships. An interactive fitting environment for Fusion 360 and a step-by-step guide are available in the project repository. A hexagonal recess matching the implanted standoff provided a keyed, repeatable mounting orientation across sessions. After printing, IR mirrors were inserted into their dedicated sockets, and IR LEDs were wired into the built-in channels and secured with adhesive, then coated with a thin layer of epoxy resin. Comprehensive virtual fitting guide, printing specifications, and project files are available in Supplementary Methods and associated repository.

### Alternative modular headmount solution

In rapidly recovering animals (e.g., mice), normal behavior is resumed before a custom single-piece headmount can be printed (which may require >10 hours). To bridge this gap, we used a modular headmount kit consisting of interlocking, ball-jointed camera holders connected to the slotted anchor by adjustable support arms (Fig. 1c). Support arms spanning a range of lengths and angles were printed in advance. Following the photogrammetry session, fitting was performed under anesthesia: the modular headmount was assembled on the hexagonal anchor, and cameras were installed on the support arms and aimed at the eyes using live video feedback under IR illumination. Once alignment was achieved, ball-joint positions were fixed with UV-curing adhesive. The headmount was then removed for wiring and epoxy coating, as in the single-piece design.

### Video acquisition

Eye videos were acquired using miniature IR-sensitive board cameras (Arducam OV5647 “NoIR” modules) installed on the custom headmount. Each camera was fitted with a manual-focus M7 lens (3.7 mm focal length) and a custom IR-pass filter (>850 nm) to isolate pupil reflections. Cameras viewed the eyes indirectly via 940-nm reflective mirrors, which remain visually transparent to the animal. This mirror-based optical path enabled stable, close-range imaging (working distance ∼2-4 cm) while keeping the hardware outside the animal’s direct field of view. Camera configuration and acquisition were controlled using custom Python code running on each Raspberry Pi via SSH, typically running at 640×480 px resolution and 60 frames-per-second. The software generated a digital frame-acquisition event for every captured frame, which was sent to the Open-Ephys system for temporal alignment across data streams. Illumination was provided by two 940-nm IR LEDs per camera, powered by a Cyclops LED driver. The driver was programmed to briefly suspend LED power for 32ms at the end of each minute and transmitted a LED-ON event signal to an Open-Ephys acquisition board for offline synchronization between eye videos, accelerometer data and external behavioral recordings. Recording sessions (usually 3-4 hours) were broken into 30-minute blocks.

### Video stability assessment

Camera stability was quantified by tracking frame-to-frame shifts of a fixed region of interest (ROI) in the eye video. For each block and eye, a static ROI was manually selected on a stable feature of the image (headmount feature when available or stable mirror edges or prominent stationary scales in *P. vitticeps*). The ROI was extracted for all frames, and a normalized 2-D cross-correlation was computed between the ROI of the current frame and that of the first frame in the sequence. The location of the correlation peak was taken as the best alignment between frames, and its displacement relative to the reference peak (frame 0) was recorded for each frame. Per-frame jitter was computed as the Euclidean distance of this displacement, which was subsequently converted to physical units using an individually calibrated pixel-to-millimeter scale derived from anatomical landmarks. These displacements were aggregated across stable periods to yield a per-camera jitter distribution.

### Behavioral arena and external video acquisition

Experiments were conducted in a custom behavioral arena (70 x 100 x 45 cm) equipped with four overhead and side-view video cameras providing synchronized coverage of the full arena. External behavioral videos were recorded continuously at 60 frames-per-second. For recordings involving head-movement analysis, animals also carried an Intan RHD-2132 headstage, and accelerometer measurements from the onboard sensor were acquired directly by the Open Ephys system. External video streams and accelerometer signals were used to annotate behavioral state (active vs. quiet) and to verify the correspondence between eye movements and whole-body activity. Mice were placed within their home cage inside the arena for recordings.

### Data Synchronization

Digital frame-acquisition signals from the left and right eye cameras, LED-state signals from the Cyclops driver, and external video events were recorded concurrently using an Open Ephys acquisition board. For each recording block, these digital event logs were parsed to construct an initial mapping between Open Ephys timestamps and eye-camera frame indices, external video frames and accelerometer samples (already recorded directly in Open Ephys). Synchronization was visually validated using the 32-ms LED-off events programmed into the Cyclops driver. All LED-off events were automatically detected across modalities and compared. When discrepancies occurred, a small user-applied temporal correction was made via a Python interface. After verification, a per-block synchronization table was generated containing frame-level correspondences between eye videos, external behavioral videos, and accelerometer data. All analyses used these synchronized time bases (Fig. S6, Video S1).

### Video analysis and eye angle reconstruction

Pupil positions were first estimated frame-by-frame using DeepLabCut, and frames with low-confidence detections were excluded. For each valid frame, a least-squares ellipse was fitted to the pupil boundary, yielding its center and major/minor axes. Angular eye position was then computed using a geometric method derived from Kerr et al. (2013), which infers azimuth (φ) and elevation (θ) from two measurable image quantities: the pupil-ellipse eccentricity (minor/major axis ratio) and the pupil-center displacement relative to the condition where the eye is pointing directly along the camera axis. In this framework, changes in ellipse eccentricity determine a depth-like scaling factor that relates pupil-center offsets in the image to rotations of the eye, and φ and θ are obtained by converting these normalized offsets into angular displacements using inverse-trigonometric relationships [25]. Because animals rarely look exactly at the camera, the “straight-ahead” reference position required for this computation was determined geometrically. The two IR LEDs mounted near each camera produce reliable reflections in the pupil. By annotating frames in which both reflections were visible, we identified the pupil location corresponding to the camera axis for each eye and used this as the zero-angle reference for subsequent projection. To quantify the accuracy of this reconstruction procedure under our optical geometry, we evaluated it using a Blender-generated synthetic dataset in which a 3D eye was rotated systematically through known azimuth-elevation combinations (-60 to +60) (Blender file available in the project repository). Applying the full ellipse-fitting and projection pipeline to the rendered images yielded angular errors that increased with eccentricity, ranging from ∼1 near the center to ∼5 at the maximal eccentricities reached by *P. vitticeps* during behavior when taking camera tilt into account (Fig. S1). Before plotting, φ and θ traces were recentered from the camera-centric 0,0 alignment to anatomically comparable resting positions by subtracting their block-wise medians. Pupil diameter (mm) was computed from the ellipse major axis using the same pixel calibration value used for camera jitter quantification.

### Head Movement Detection and Behavioral state segmentation

Where mentioned, head-movement activity was quantified from the three-axis accelerometer embedded in the Intan RHD2132 headstage, which was attached to the animal’s head via an implanted connector. Raw acceleration traces were processed using custom MATLAB code to identify movement events by first estimating the noise distribution from low-variance segments of the filtered signal (selected using kurtosis-based criteria) and then classifying timepoints as movement when the acceleration magnitude exceeds the noise baseline by several standard deviations. The resulting time-stamped movement events were imported into Python and used to annotate saccadic events (see below). Movement events were also converted into a continuous behavioral annotation by applying a rolling-window analysis (10-s window, 1-s step). For each window, the mean movement magnitude was computed, and windows were labeled as active when the mean exceeded an adaptively chosen threshold (user verified per recording) and quiet otherwise. Consecutive windows of matching labels were merged into contiguous behavioral epochs, used for all state-dependent analyses in the manuscript.

### Saccade detection and segmentation

Eye-movement events were detected separately for each eye by computing frame-to-frame changes in reconstructed eye angles (φ, θ) and identifying periods in which the instantaneous angular speed exceeded a fixed threshold (50 deg/second). Consecutive suprathreshold frames were grouped into candidate saccades. Because some events contained multiple direction changes, each candidate event was subdivided whenever the instantaneous movement direction shifted by more than 90 deg, and only segments with a minimum duration (2 frames) and a net angular displacement >0.5 deg were retained. For each qualified saccade segment, onset and offset times and net angular displacement were extracted. When head-movement information was available, each saccade was labeled as “with head movement” if its duration overlapped any accelerometer-defined movement event (as described above), and “without head movement” otherwise. Binocular classification was performed by matching left and right eye saccades based on their onset time. Pairs whose onsets differed by <34 ms were classified as synchronized binocular saccades, and all remaining events were treated as monocular. All subsequent analyses of saccade amplitude, direction, dynamics, and pupil behavior used these classified sets. For Fig. 2f, the ipsilateral peak for each event was taken from the detected saccade, while the contralateral peak is the maximal instantaneous speed within ±51 ms of onset (irrespective of contralateral detection). Because onset-based pairing can undercount binocular saccades when temporal drift occurs between eye videos within the 1-minute verified epochs, we implemented an additional step to define a conservative set of verified monocular events. For each non-synchronized (candidate monocular) saccade detected in one eye (the “ipsilateral” eye), we examined the continuous ocular speed trace of the contralateral eye directly, irrespective of whether a contralateral saccade was detected by the segmentation algorithm. We computed the contralateral peak instantaneous angular speed within a symmetric temporal window around the ipsilateral onset (±51 ms, i.e., ±3 frames at 60 Hz). If the contralateral peak speed remained below the saccade threshold (0.8 deg/frame, ≈50 deg/s) throughout this window, the event was classified as verified. If contralateral peak speed exceeded threshold, the event was tagged as unverified. Events for which contralateral traces were unavailable in the verification window were excluded from the verified monocular set (Video S2). All reported “monocular” fractions in the analysis were computed using this verified subset. To compute Inter-saccade-interval histograms (Fig. 3d), ISI values were collected within each recording and probabilities were calculated for each animal over the full ISI range (10-40,000 ms), then applied uniformly to both panels.

### Accelerometer-based tests of head movement effects on saccade amplitude and rate

To test the relationship between head movements, saccade amplitude, and saccade rate, we used accelerometer recordings to derive two labels: (i) a binary head-movement tag for each saccade (based on whether the saccade overlapped suprathreshold accelerometer activity), and (ii) accelerometer-defined behavioral state (active vs quiet) by binning the recording into non-overlapping 1-s epochs and thresholding movement magnitude.

#### Amplitude

For each animal with accelerometer data (n=4), saccades were pooled across blocks and split by the head-movement tag; we computed Δamplitude as the difference in mean saccade amplitude between head-movement and no-movement saccades, and summarized the effect as the mean Δamplitude across animals.

#### Rate

Using the accelerometer-defined active/quiet bins, we computed state-specific saccade rates as total onsets divided by total time in each state, yielding Δrate = rate_active − rate_quiet, and summarized the effect as the mean Δrate across animals. For both analyses, significance was assessed with within-animal Monte-Carlo permutation tests (randomly shuffling head-movement labels across saccades, or state labels across time bins while preserving counts) to generate null distributions of the mean effect, from which two-sided p-values were computed.

### Head movement and saccade amplitude

To test whether head movement was associated with saccade amplitude, saccades were pooled across blocks within each animal and split by a binary head-movement label. For each animal (n=4 animals with accelerometer measurements) we computed the difference in mean amplitude between conditions, and summarized the effect as the mean Δamplitude across animals. Significance was assessed with a within-animal Monte-Carlo permutation test. Head-movement labels were randomly reassigned across saccades to generate a null distribution of mean Δamplitude values, from which a two-sided p-value was computed.

### Behavioral state and saccade rate

To test whether behavioral state predicted saccade rate, active / quiet annotations were converted into non-overlapping 1-s time bins, and saccade onsets were counted per-bin. For each animal, rates were computed as total saccades divided by total annotated time in each state, yielding Δrate (rate_active - rate_quiet), and the population effect as the mean Δrate of this across animals. Significance was assessed by Monte-Carlo permutation of state labels across time bins within each animal (preserving the number of active/quiet bins) to generate a null distribution of mean Δrate values, from which a two-sided p-value was computed.

### Pupil data analysis

Pupil diameter (mm) was derived from the fitted ellipse parameters in each eye-tracking frame by converting the major axis from pixels using the per-eye pixel calibration described above. Frames with low DeepLabCut likelihood or unstable ellipse fits were excluded. To characterize state-dependent changes in pupil size, pupil traces were aligned to behavioral annotations, and distributions were computed separately for active and quiet periods. For each animal, pupil diameters from both eyes were combined, clipped to remove extreme outliers, and z-scored relative to the animal’s pooled distribution. State-dependent effects were quantified as the difference between the active and quiet z-scored histograms (Active - Quiet), allowing comparison across animals despite inter-individual differences in absolute pupil size. This reported, for each normalized diameter bin, whether that range is sampled more frequently during active or quiet behavior (Fig. 3f). To assess whether these differences exceeded chance, we performed a permutation test separately for each animal by randomly shuffling the active/quiet labels within each block and eye while keeping the pupil time series fixed. For each shuffled dataset, we recomputed the difference in median pupil diameter between quiet and active epochs and used the resulting null distribution to obtain a two-sided p-value for the observed median difference. Inter-eye pupil coordination was quantified from the z-scored traces by computing rolling Pearson correlations between left and right pupil diameter in 10 second windows, requiring at least one third of samples per window. Each correlation value was labeled by behavioral state and classified as positively correlated (r > 0.2), weakly coupled (-0.2 ≤ r ≤ 0.2), or anti-correlated (r < -0.2). For each animal, the fraction of time spent in each category was calculated separately for all, active, and quiet epochs and visualized as stacked bar plots.

### Orientation and directional analysis

To characterize the directional structure of eye movements, each detected subsaccade was represented by its net angular displacement in azimuth-elevation space (Δφ, Δθ). For visualization, all saccades were translated so that their starting positions lay at the origin, and their endpoints were plotted in this common coordinate frame. This produced an egocentric map of saccade directions that is independent of the animal’s instantaneous eye position. For saccade target heatmaps (Fig. 2h), densities were estimated with a 2D Gaussian kernel on a ±20 degree grid. For each animal, saccade events were histogrammed and a main movement axis was identified by scanning candidate orientations (0-180) and selecting the axis with maximum saccadic events (Fig. 2i). To determine the movement bias for saccades along this axis, on-axis events were defined as within ± 45° degrees of the main movement axis. Monocular and binocular saccades were analyzed separately, and the bias for each animal was displayed for concurrent and monocular events (Fig. 2j).

## Supporting information

Document S1. Supplementary methods

Document S2. Figures S1–S6

Video S1. Synchronized video example: 55 seconds of synchronized video output for (left-to-right) right eye, top-down arena camera, and left eye. Videos are shown raw, with eye data traces showing eye angle and pupil diameter below. red=right eye, blue=left eye.

Video S2. Monocular saccades example: A collection of synchronized videos for events detected as monocular saccades. Green dot denotes saccade detection. Slowed down to 20% speed.

