## Supplementary Methods for "A personalized eye-tracking system reveals complex state-dependent visual acquisition in lizards"

### Detailed protocol

#### Cranial Anchor Implantation and Photogrammetry

This procedure describes a minimally invasive protocol for surgically implanting a lightweight cranial anchor for the eye-tracking headstage onto the skull of *P. vitticeps*. The same overall approach was used in turtles and mice with minor adjustments to anchor size and fixation, but the core steps were identical.

Before surgery, animals received analgesic, anti-inflammatory, and antibiotic treatment to minimize discomfort and infection risk. We administered Meloxicam or Carprofen at 0.2 mg/kg (diluted to 1 mg/ml), Baytril (enrofloxacin) at 5.0 mg/kg (diluted to 1 mg/ml), and Dexamethasone at 0.5 mg/kg. All surgical materials were prepared in advance and instruments were thoroughly sterilized. The anchor location was chosen along the midline of the dorsal skull in regions where the bone is sufficiently thick to support shallow drilling and a strong adhesive bond.

Anesthesia was induced in an isoflurane induction box and the animal was then intubated and ventilated with 4% isoflurane (AWS 100, Hallwell EMC). The animal was placed in a stereotactic apparatus (RWD 68409) on a heating pad set to 35 deg C to maintain body temperature. Ophthalmic ointment (Duratears) was applied to both eyes to prevent drying. The skin overlying the dorsal skull was disinfected with 10% povidone iodine, rinsed with double-distilled water, and coated with a thin layer of 2% lidocaine ointment. We exposed only the portion of the skull where no muscle lies between skin and bone: a blunt tool was used to palpate the transition between muscle and bare skull, and a marker pen was used to delineate the skin area to be removed. This patch of skin was then excised carefully with a scalpel so as not to damage the underlying bone. Residual soft tissue was cleaned using a micro curette, and remaining fragments were dissolved with 30% hydrogen peroxide until a clean, dry bony surface was visible. Where applicable (*T. elegance* and *P. vitticeps,* but not *M. musculus*), stainless-steel screws were added to strengthen anchor fixation. In these cases, 2-4 M1 pilot holes were drilled into stable, visibly thick areas of the skull, taking care not to breach the cranial cavity. Short, sterilized M1 screws were inserted into these holes and tightened just enough to verify rigid stability without applying excessive force to the bone. The exposed skull was then etched using Dentaflux (gel de Acido grabador azul) and coated with Evatric Bond to create a uniform adhesive surface. The anchor itself (M3 brass hexagonal standoff weighing 0.6 grams for lizards and turtles, smaller M2 plastic hexagonal standoffs weighing 0.15 grams for mice) was prepared by drilling a circular small indent into its circumference near the base to provide mechanical purchase for the adhesive. Using the stereotaxic arm, the anchor was positioned over the prepared midline site and adjusted so that it was level and correctly oriented to support the future camera geometry. UV-curing glue (Transbond XT, 3M) and dental cement (Coral Fix) were applied to connect the etched skull surface, the reinforcement screws (when present), and the anchor base into a single rigid construct. The dental cement was allowed to dry completely before any further handling.

We proceeded to photogrammetry while the animal was still under anesthesia. The goal of this step was to obtain a precise 3D model of the head, including the eyes and the implanted anchor, for subsequent virtual headstage fitting. We typically used a Sony RX100-II camera, but any high-definition camera that records acquisition-angle metadata can be used. A better camera and sharper images improve the resulting 3D model and texture quality. Lighting conditions did not need to be spatially uniform, but they were kept stable during acquisition. Changes in illumination between images degrade reconstruction quality. For each animal, we collected 40-60 high-quality images covering all relevant angles and features of the head (including the anchor and both eyes), ensuring that each image was in focus and free of motion blur. For a precise reconstruction, the animal must remain as still as possible. If movement occurred before sufficient images were acquired, the partial dataset was discarded and imaging restarted. Uniform and/or reflective backgrounds such as stainless-steel tables, plain white walls and reflective objects were avoided, as the reduced feature richness introduced reconstruction artifacts. To reduce specular reflections on the metallic anchor we applied a small piece of masking tape with a scribbled X to the exposed socket, which dramatically improved the 3D reproduction (gleaming surfaces are harder to reconstruct due to varying reflections at different angles). The exact Meshroom settings used for reconstruction and an example dataset are provided in the associated project file in the project repository.

After photogrammetry, animals were allowed to recover in a warmed enclosure until they resumed normal behavior. Recovery typically required 12-24 hours for reptiles, and approximately 60 minutes for mice. Postoperative care included close monitoring during recovery and administration of analgesics and antibiotics at the same dosages as in the preoperative phase for five consecutive days. Animal weight was monitored weekly as an indicator of stress. Individuals whose weight fell below 80% of their pre-surgical baseline were euthanized according to institutional guidelines.

**Headstage Design and Virtual Fitting**

The photogrammetry-derived 3D mesh served as the anatomical basis for individualized headstage design. We imported the mesh into Autodesk Fusion 360 and rescaled it to true physical dimensions by matching the digital representation of the implanted hexagonal standoff to its measured real-world size. The mesh was then cleaned to remove background details and lightly decimated to facilitate interactive manipulation while maintaining details in critical regions like the eyes and anchor. Predefined modular components including the camera holders, mirror holders, and anchor interface, were loaded into the workspace as independent bodies. To form the headstage, these modules were positioned separately in their proper locations relative to the animal mesh, anchor and eyes, then joined together to create a single-piece rigid model. First, the anchor interface was positioned in the only possible orientation, defined by the anchor’s actual position on the 3D animal reconstruction. Next, the camera-mirror assemblies were oriented so that each camera’s optical axis intersected the eye via the reflective surface of its corresponding mirror (the camera-mirror geometry is predefined by the interaction of the camera module and the mirror holder, ensuring reproducible angles even when a mirror breaks and is replaced). Camera inclination was defined relative to a virtual horizontal axis connecting the two eye centers, and adjusted so that the cameras could maintain focus across the target working distance (typically 1.5-4 cm in *P. vitticeps*, and slightly longer (4-6 cm) with the extender module used in *T. s. elegans)* with inclinations ranging between 20-30 degrees (cameras were typically above the eyes, facing outwards and downwards at the mirror (Fig. 1)). Once alignment was finalized, the modular elements were fused into a single rigid body using either a loft-based procedure or the generative design option in Fusion 360. This single-piece headstage was then exported and prepared for printing using Formlabs's PreForm software. Because the headstage must preserve precise geometric relationships, functional surfaces such as the internal faces of the camera holders and the hexagonal recess were kept free of support structures by carefully choosing printing orientation (support touchpoints leave small bumps or indentations on the finished 3D print even after removal and cleaning). The resulting file was printed in a Formlabs model 2B using Grey V4 resin at 25 micron resolution, producing headstages that were mechanically robust, dimensionally consistent, and sufficiently lightweight for use in freely behaving small animals.

To facilitate adoption, we include a Fusion 360 file with adjustable components as part of the project repository ('Virtual_Fitting_Main.f3d'), and include a step-by-step guide for its usage.

**Camera and Optical System**

The eye-tracking cameras were based on commercially available Arducam NoIR modules (OV5647 sensor), which were mechanically and optically modified to allow infrared imaging at short working distances. Each camera was first stripped of its stock plastic lens assembly to expose the bare sensor and accommodate a custom M7 threaded lens mount. A 3.7-mm manual-focus M7 lens (QILENS) was prepared by attaching a custom cut infrared-pass filter (>850 nm) directly to the front rim of the lens. Filters were cut as small circular discs (approximately 2.5 mm in diameter for a lens rim of ~2mm) using a laser cutter. To affix the filter without introducing optical artifacts, the lens and filter were held together under uniform pressure using a small custom-built vice with soft-foam contact faces, and a single micro-drop of cyanoacrylate was applied at the perimeter. The modified lens was inserted into the glued-on M7 threaded mount on the camera board. Care was taken to avoid both glue vapor contamination and mechanical stress on the sensor, as any residue on the sensor surface would markedly degrade image quality. Each camera holder module of the headstage provided two angled slots adjacent to the camera holders for mounting illumination LEDs. Into these, two 940 nm infrared LEDs (Vishay VSMB2943GX01) were positioned and permanently affixed using UV-curing adhesive or cyanoacrylate. LED orientation and slot geometry were predefined to produce uniform illumination of the eye through the IR mirror without producing glare. The four LEDs of a headstage were wired in-line and connected to a Cyclops LED driver, which provided stable and smoothly adjustable illumination. The driver's current limit was set to 100 mA per LED, 400 mA maximum. The cameras were aimed at the eyes using small 8x8 mm infrared-reflective mirrors (940 nm), affixed to holders with dedicated grooves to guide mirror placement, which fit within a slot on each camera module to support the mirror in front of the camera. Since this is the most at-risk part of the headstage as the mirror protrudes from the head boundary, we made the mirror holder easily replaceable. The mirror-camera geometry is hard-coded in the modular design, preserving the angle between camera, mirror, and eye when replaced. During each session, camera focus was adjusted manually by threading the lens along the M7 mount while viewing a live feed on an HDMI-connected display attached to each Raspberry Pi. Final focus was set to produce a crisp pupil contour. To prevent mechanical load on the animal, camera and LED cables were routed through a counterbalanced pulley system. A thin fishing line passed through two freely rotating plastic rollers above the arena, with an adjustable water-filled counterweight on one end and a metal clip holding the camera cables on the other. The counterweight was tuned for each session such that the headstage was neutrally balanced at the center of the arena. Cables were relatively long (2 meters from pulley) to minimize shear forces on the animal head during behavior.

**Raspberry Pi Video Acquisition and Synchronization**

Eye videos were acquired using Raspberry Pi 4B computers running custom Python software based on libcamera and the picamera2 module. Each Raspberry Pi controlled a single camera, encoded videos in H.264, and sent a TTL pulse for every acquired frame to the Open Ephys acquisition system. All eye-video data were aligned directly to the Open Ephys clock using the recorded frame-acquisition TTLs. During development of the synchronization pipeline, we examined mean brightness values across frames and compared them to the Open Ephys LED-OFF events (brief 32-ms LED power suspensions occurring once per minute). Because these LED-OFF periods are visible in both eye videos and recorded as TTLs, we expected a perfect temporal match between the drop in video brightness and the corresponding digital event. We consistently observed that the digital LED-OFF event did not perfectly coincide with the synchronized eye video frames, which likely reflects a fixed sensor-to-TTL latency in the Raspberry Pi acquisition stack, though we did not identify the exact source. Importantly, the offset was stable within each recording block yet differed between blocks and between Raspberry Pis. To correct this constant latency, we developed a custom Python GUI that jointly displays the eye-video brightness trace and the LED-OFF TTL series. For each block, the user identifies the corresponding LED-OFF frame based on brightness values and applies a single temporal shift to align the video stream with the Open Ephys event. Once a correction was applied, all other LED-OFF events were used to verify block-wide alignment.

Pupil positions were obtained using DeepLabCut (DLC) with eight points along the pupil boundary, and two marking the rostral and caudal tear ducts. Networks were trained using the standard ResNet-50 backbone. Frames were retained only when all pupil-contour labels exceeded a likelihood threshold of 0.998. Additional frame rejection was performed downstream: frames with abnormally small pupil diameters (indicative of partial eyelid closure) were removed automatically, and segments with erratic or unstable pupil-diameter behavior were flagged and manually reviewed using a custom inspection tool available in the project repository.

For each valid frame, the pupil contour was fit with a direct least-squares ellipse[43], yielding the pupil center, axis lengths, and ellipse rotation angle. These parameters served as the basis for pupil-diameter estimates and for reconstruction of eye orientation. Angular gaze coordinates ($\varphi$, $\theta$) were computed using a geometric method which infers azimuth and elevation from pupil-center displacement relative to a reference position and the ellipse minor/major axis ratio[15]. A zero-angle reference was defined separately for each eye and recording block. Frames in which both LED reflections were simultaneously visible in the pupil were identified using a dedicated GUI. Since LED-camera geometry is known, such frames provided the camera axis coordinate in frame - the gaze angle was defined as 0,0 when the pupil center is aligned with this coordinate and the minor/major axis ratio approaches 1.

Reconstruction accuracy was evaluated using a synthetic dataset generated in Blender, where a 3D eye model was rotated through a grid of known azimuth-elevation values (-60 to + 60). The same ellipse-fitting and reconstruction pipeline was applied to the rendered images. Errors were smallest near the optical axis and increased toward the extremes, but remained within the span of ± 5 degrees over the eccentricity range used by *P. vitticeps* (typically within ±35 degrees, Fig.S1) Before analysis, φ and θ traces were re-centered by subtracting the block-wise median resting position to improve comparability across sessions and animals.

**Saccade Detection and Segmentation**

Saccades were detected independently for each eye using frame-to-frame changes in reconstructed gaze angles. Angular speed was computed using a first-order temporal derivative applied separately to φ and θ without temporal smoothing, except in plots where interpolation was explicitly noted (Fig. 2c,d). A saccade was defined as any continuous sequence of frames in which the 2D angular speed exceeded 50°/s. Candidate saccades often contained multiple directional components. To isolate physiologically meaningful submovements, events were subdivided whenever the instantaneous movement direction changed by more than 90°. This threshold was chosen because a 90° turn within a ~17 ms window was empirically associated with a distinct, newly initiated targeting movement rather than the curvature of a single ballistic saccade. Sub-events shorter than two frames or with a net displacement below 0.5° were discarded. Onset was defined as the first frame exceeding the speed threshold and offset as the first frame after peak speed at which angular speed dropped back below threshold. For binocular classification, left- and right-eye saccades were matched based on onset time. Events whose onsets differed by less than 34 ms were considered synchronized binocular saccades; all others were treated as monocular. When raw accelerometer data were available, each event was additionally labeled as "with head movement" if its duration overlapped any accelerometer-defined head-motion event. This distinction allowed separation of eye-driven gaze shifts from those occurring during coupled eye–head movements. All subsequent analyses of amplitude, speed scaling, directionality, coupling and state-dependent dynamics used these segmented and classified saccade events.

**Pupil Diameter Analysis**

Pupil diameter was computed from the major axis of the fitted ellipse in each valid frame. Pixel-to-millimeter conversion was performed separately for each block using a visible anatomical reference when available (e.g., a stable scale or tear-duct distance where steady annotations were available), and manually normalized across blocks of the same animal when required. Because absolute diameter comparisons across animals were unreliable due to geometry and focus differences, all cross-animal analyses were performed on normalized (z-scored) diameter values within each animal. To examine behavioral modulation of pupil size, diameter traces were aligned to behavioral-state annotations and grouped into Quiet and Active epochs. Distributions were constructed from all valid samples within each state, combining left and right eyes per animal. For each animal, values were z-scored relative to the pooled distribution across both states, and Active–Quiet difference curves were computed in matched diameter bins to visualize shifts in sampling probability. Inter-ocular coordination was quantified using rolling Pearson correlations (10-second windows, requiring at least one-third of samples). Correlation values were categorized as positively correlated ($r > 0.2$), weakly coupled ($-0.2 \leq r \leq 0.2$), or anti-correlated ($r < -0.2$). Time spent in each category was computed separately for Active, Quiet, and all epochs. These metrics were used directly for the state-dependent coupling analyses (Fig. 3, S4).

**Behavioral State Segmentation**

Behavioral state was quantified using the three-axis accelerometer embedded in the Intan RHD-2132 headstage, recorded directly by the Open Ephys system and synchronized with the eye videos as described above. Raw acceleration traces were first filtered and then processed to estimate baseline noise levels. Low-variance segments were identified using kurtosis-based criteria and the standard deviation of these segments served as an estimate of sensor noise. Accelerometer samples were classified as movement when the magnitude of the filtered signal exceeded this noise baseline by a user-verified factor, fixed per accelerometer unit. Movement events were then converted into a continuous state annotation using a rolling analysis. For each 10-s window (advanced in 1-s steps), the mean movement magnitude was computed. Windows whose mean exceeded a user-defined threshold (per accelerometer) were labeled Active, and all others labeled Quiet. External behavioral videos taken from above the animal were inspected to validate accelerometer-based labels. These annotations were used in all state-dependent analyses of saccade rate, pupil diameter, and inter-ocular pupil coupling.

**Directional Eye Movement Analyses**

Directional structure of eye movements was quantified from the net angular displacement of each segmented subsaccade. For every saccade event, the start and end gaze positions in azimuth and elevation (φ, θ) were extracted, and the displacement vector (Δφ, Δθ) was computed. To visualize directionality independent of absolute gaze position, all saccades were translated so their starting point lay at the origin, yielding an eye-centered endpoint map for each eye and animal. Inspection of these endpoint distributions revealed a clear dominant movement axis for synchronized binocular saccades, whereas monocular saccades showed a broader and more isotropic pattern. This dominant axis was then used as a reference to align data across animals: for each eye, the saccade cloud was rotated such that the dominant axis lay at 0°/180°. This procedure accounted for small differences in headstage orientation and camera alignment, enabling cross-animal comparisons in a common coordinate frame. Directional bias was quantified relative to this aligned axis. For each animal, we computed:

$$bias = \frac{P(along - axis) \; - \; P(perpendicular)}{P(along - axis) \; + \; P(perpendicular)}$$

where “along-axis’’ saccades were defined as those whose displacement direction lay within ±45° of the dominant axis, and “perpendicular’’ saccades were defined relative to the orthogonal axis. Bias values were computed separately for synchronized and monocular saccades, using the same rotation for both categories.

Because camera coordinates vary with head orientation and we did not track world-referenced head pose, all analyses were performed strictly in this eye-centered coordinate system without interpretation in external spatial units. Block-wise medians were subtracted from φ and θ before analysis to correct for small differences in resting eye position across sessions.

**Video Stability Assessment and Jitter Rejection**

Video stability was quantified for each eye and recording block by tracking frame-to-frame motion of a fixed region of interest (ROI) in the eye image. A stationary feature was manually selected for each block, typically a part of the headstage, a stable mirror edge, or a patch of non-moving facial scales. For every frame, this ROI was compared to the reference ROI from the first frame using normalized 2D cross-correlation, and the location of the correlation peak provided the best alignment estimate. The Euclidean distance between the peak location in frame t and frame 0 defined the per-frame jitter. Pixel distances were converted to physical units using the per-eye pixel calibration determined during pupil-size conversion. Jitter values were monitored throughout each block to detect mirror flex, cable tug, or brief mechanical disturbances. Frames with displacement values exceeding 30 pixels were excluded from analysis (this typically occurred when the animal contacted the arena walls and caused temporary deflections of the mirror holder. In many such cases, the corresponding DLC pupil detections were already invalid, so these frames were removed redundantly by both procedures. Data were retained once displacement returned to the typical range (0-10 pixels). Blocks were not discarded solely on the basis of jitter unless the mirror was physically damaged, which produced a persistent loss of stability. In such cases, the recording was terminated, the mirror-holder module was replaced, and a new recording block was initiated. No entire animal was excluded due to jitter-related issues. Across the full dataset, 368,470 frames out of 2,128,904 total jitter-quantified frames were rejected (~17%).

**Behavioral Recording Arena**

Eye-tracking experiments were conducted in the PreyTouch behavioral arena described in detail in Eyal et al., 2024. In brief, the arena measured 70 × 100 × 45 cm and was uniformly illuminated using overhead white LEDs powered by a constant 12 V, 5 A supply to avoid flicker. Four synchronized high-speed arena cameras (FLIR monochrome, 60 fps) were positioned above and at the sides of the enclosure to provide continuous coverage of the animal's full body posture and locomotion. These cameras delivered frame-acquisition TTLs directly to the Open Ephys system, enabling alignment with eye-camera recordings and accelerometer data. For experiments requiring head-movement annotation, animals carried an Intan RHD-2132 headstage, and the three-axis accelerometer embedded in the unit was recorded synchronously through the same Open Ephys acquisition pipeline. A complete description of the arena construction, camera synchronization hardware, illumination uniformity, and optional peripheral modules (reward delivery, motion sensors, thermocontrol) can be found in Eyal et al. (2024), and all relevant details for replicating the core setup are provided there.
