## Supplementary Figures for "A personalized eye-tracking system reveals complex state-dependent visual acquisition in lizards"

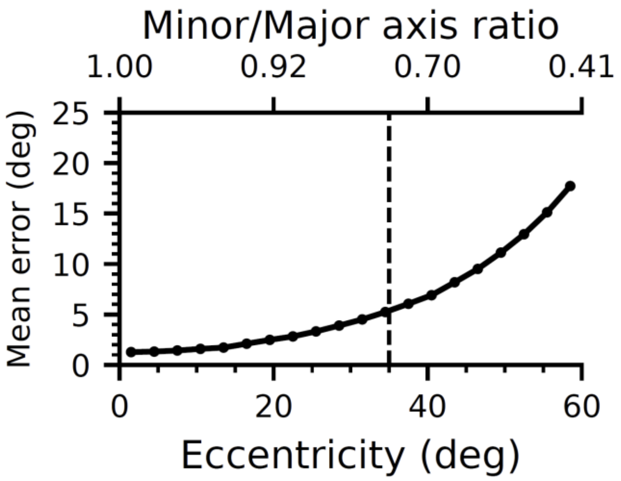


**Figure S1: Simulation based accuracy assessment** - a 3D blender animation of a rendered eye monotonously spanning the angle range of (-60,-60) to (60,60) degrees (1 degree step) was analyzed for pupil data, and results were compared with ground-truth. Mean total error of the angular gaze projection calculated as sqrt((φ - X angle)$^2$ + (θ - Y angle)$^2$) per-frame is displayed as a function of the ellipse ratio (minor/major ax) in the images, annotated via OpenCV-python (see git repository). The top x-axis shows ratio (reversed, 1 -> 0), eccentricity increases to the right. The bottom x-axis reports the representative eccentricity (deg) based on the dataset such that for each ratio bin we compute the median ground-truth radius r= sqrt(X angle$^2$ + Y angle$^2$) and use it to label the ticks. Curve is the mean error for 30 evenly spaced ratio bins. The vertical dashed line marks the ratio threshold corresponding to an extended ocular span for *P .vitticeps* (+/-35 degrees Euclidean for φ and θ (maximal radial deflection=50)).

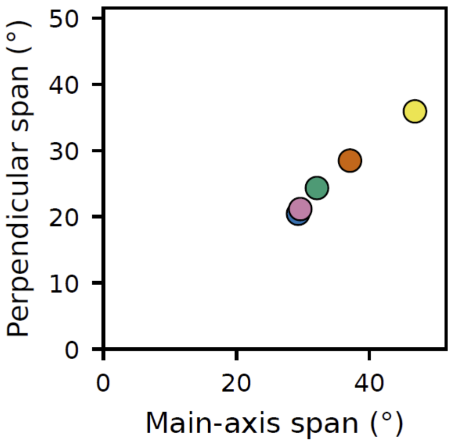


**Figure S2. Oculomotor span for each animal.** For each animal and eye, saccade directions were used to estimate an initial movement axis by identifying the peak of the bidirectional (0–180 folded) saccade-direction histogram. Gaze coordinates (φ, θ; centered relative to each eye's resting orientation) were rotated by the shortest angle needed to align this axis horizontally. The central 95% gaze span (2.5–97.5 percentiles) was computed along the rotated axes. If the axis perpendicular to the saccade-aligned axis showed a larger span, axes were swapped so that the dimension with the greater gaze range defined the main axis. Points show per-animal spans (mean across eyes). Color indicates animal identity.

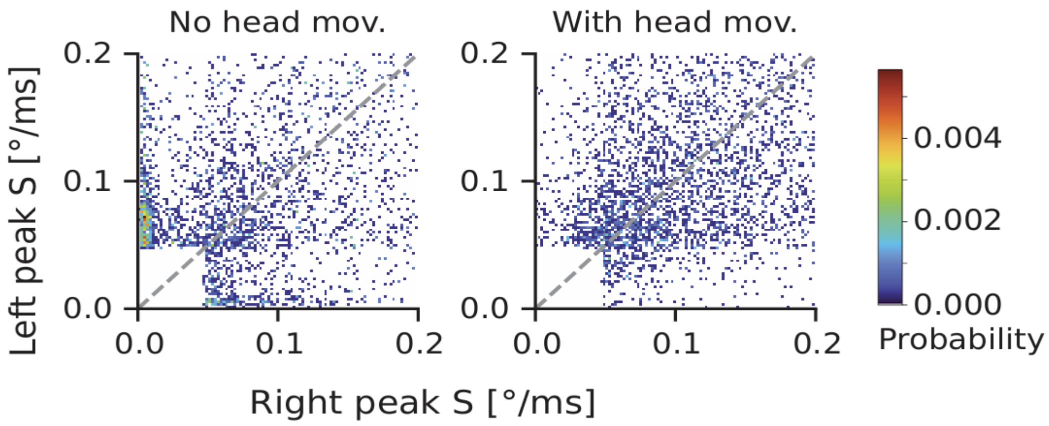


**Figure S3. Coupling of peak saccade speed between eyes, with and without head movements.** Two-dimensional histograms show the joint distribution of right- and left-eye peak saccade speed (peak S; °/ms) for saccades occurring in the absence of head movements (left) and with head movements (right). Each point contributing to the histogram corresponds to a saccade detected in one eye, paired with the maximum angular speed in the contralateral eye within a ±51 ms window around saccade onset. Candidate events were classified as binocular when a contralateral saccade onset occurred within ±60 ms of the ipsilateral onset; when binocular events were analyzed, duplicate left/right detections were merged within 60 ms to avoid double counting. Histograms were pooled between animals for equal contribution. The dashed line indicates equality between eyes. Color denotes probability density (normalized within each panel; shared color scaling)

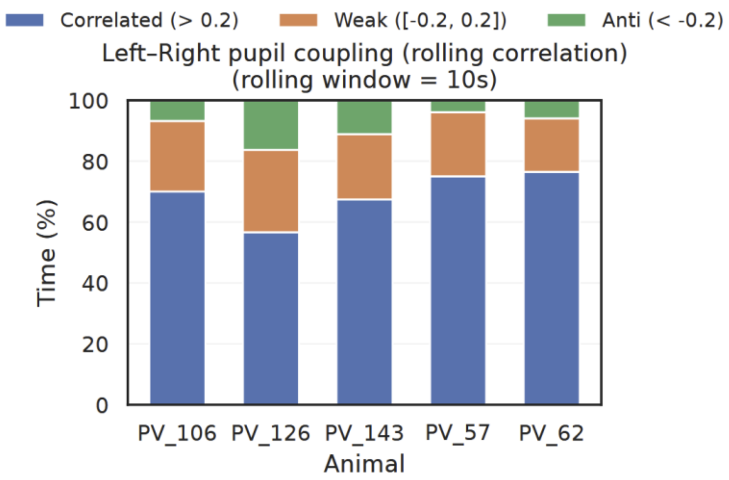


**Figure S4. correlation of pupil dynamics between eyes.** Correlated (r > 0.2, “Correlated”), weakly coupled (-0.2≤ r ≤ 0.2, “Weak”), and negatively correlated (r < -0.2 “Anti”) bilateral pupil dynamics. Fractions were computed from rolling Pearson correlations (10-s window) applied to z-scored pupil traces from all valid blocks per animal. Bars represent the time-weighted proportion of correlation states across the full dataset for each animal.

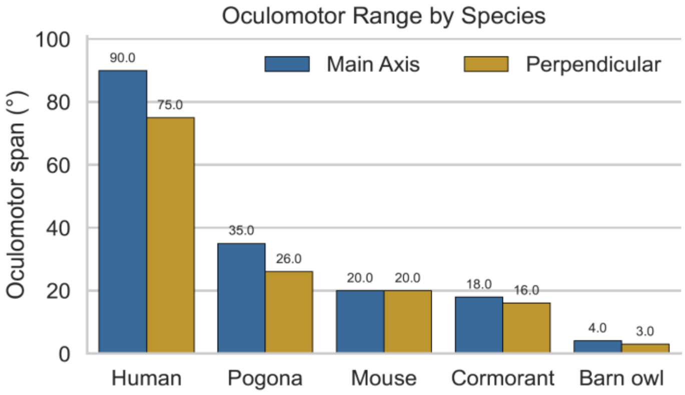


**Figure S5**. **Comparative oculomotor spans across species.** Horizontal (blue) and vertical (orange) oculomotor ranges (peak-to-peak eye-in-head rotations, in degrees) for humans, barn owl, great cormorant, mouse, and *Pogona vitticeps*. Bars show literature-derived estimates for humans (ref [78]), mice (ref [34]), barn owls (ref [84]), and great cormorants (ref [10]), alongside the spans measured in *P. vitticeps* in the present study.

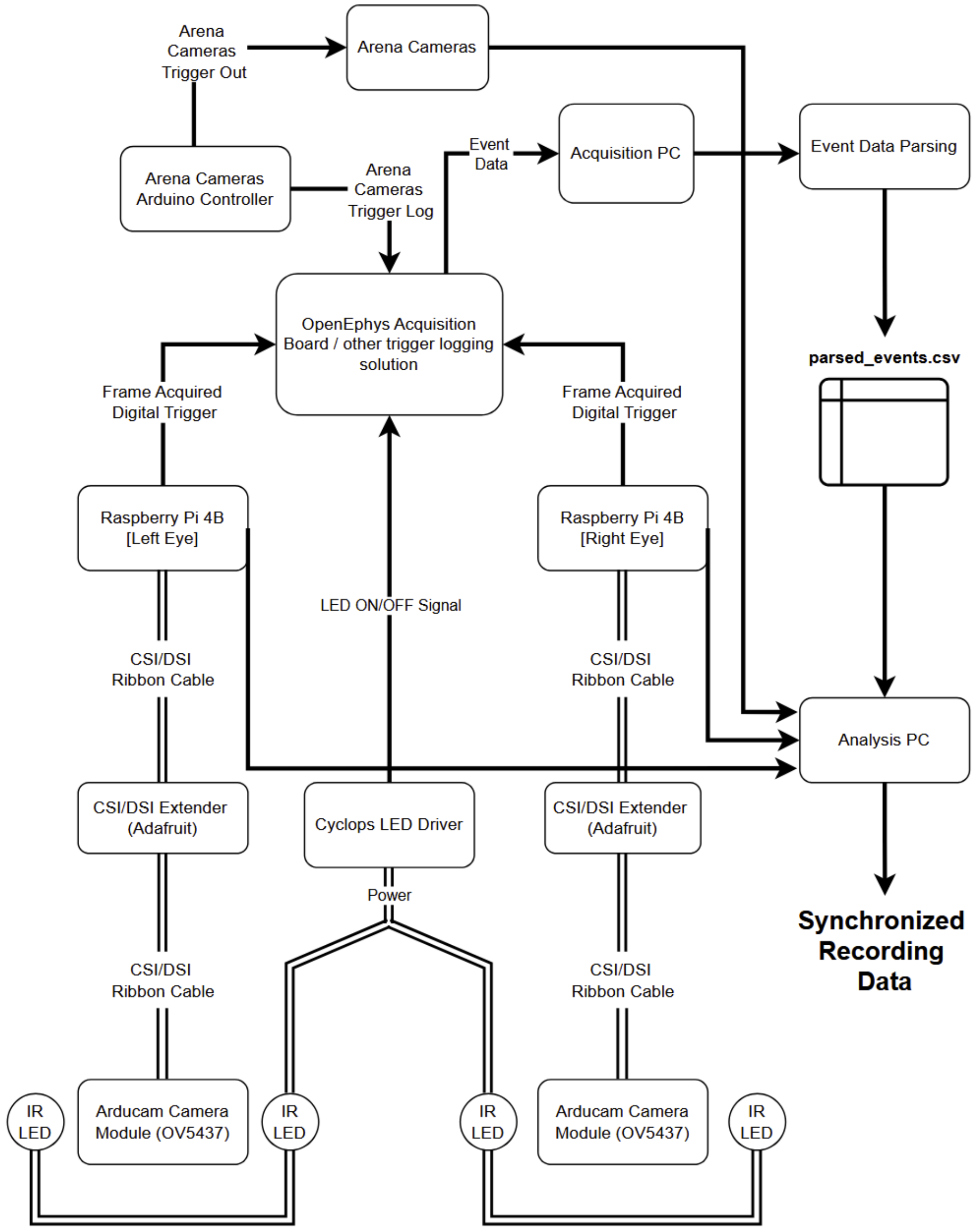


**Figure S6: Full system wiring diagram:** Double lines=physical wiring and cables to peripherals. Full arrows=data transfer via wire, cable or file transfer.
